# Combined reference-free and multi-reference approaches uncover cryptic variation underlying rapid adaptation in microbial pathogens

**DOI:** 10.1101/2022.05.16.492091

**Authors:** Anik Dutta, Bruce A. McDonald, Daniel Croll

**Author notes:** Institute of Phytopathology, Kiel University, 24118 Kiel, Germany (Anik Dutta).

## Abstract

**Background:** Microbial species often harbor substantial functional diversity driven by structural genetic variation. Rapid adaptation from such standing variation in pathogens threatens global food security and human health. Genome wide association studies (GWAS) provide a powerful approach to identify genetic variants underlying recent pathogen evolution. However, the reliance on single reference genomes and single nucleotide polymorphisms (SNPs) obscures the true extent of adaptive genetic variation. Here, we show quantitatively how a combination of multiple reference genomes and reference-free approaches captures substantially more relevant genetic variation compared to single reference mapping.

**Results:** We performed reference-genome based association mapping across 19 reference-quality genomes covering the diversity of the species. We contrasted the results with a reference-free (i.e., K-mer) approach using raw whole genome sequencing data. We assessed the relative power of these GWAS approaches in a panel of 145 strains collected across the global distribution range of the fungal wheat pathogen *Zymoseptoria tritici*. We mapped the genetic architecture of 49 life history traits including virulence, reproduction and growth in multiple stressful environments. The inclusion of additional reference genome SNP datasets provides a nearly linear increase in additional loci mapped through GWAS. Variants detected through the K-mer approach explained a higher proportion of phenotypic variation than a reference genome based approach, illustrating the benefits of including genetic variants beyond SNPs.

**Conclusions:** Our study demonstrates how the power of GWAS in microbial species can be significantly enhanced by comprehensively capturing functional genetic variation. Our approach is generalizable to a large number of microbial species and will uncover novel mechanisms driving rapid adaptation in microbial populations.

## Introduction

Rapid genetic change in microbial pathogens has led to significant damage to agricultural production as well as to human health over recent decades (Casadevall et al. 2011; Fisher et al. 2012; Figueroa et al. 2018). The rapid evolution in pathogen populations of virulence and resistance to anti-microbial drugs are key concerns in plant, animal and human health. There is an urgent need to identify the precise genetic determinants in pathogens that underlie differences in virulence and evasion of control mechanisms. Vast genomic datasets can now be exploited to retrace evolutionary pathways of pathogen adaptation. Association mapping can be used to establish relationships between genetic and phenotypic variation using field collections of pathogens (Bartoli and Roux, 2017; Sánchez-Vallet et al. 2018). The genetic variation relevant for trait evolution is often more complex than the commonly used single nucleotide polymorphisms (SNPs). Structural variants (SVs) such as insertions-deletions (indels), copy number variants, chromosomal rearrangements, inversions and duplications can also be major facilitators of microbial adaptation (Dutilh et al. 2013: Plaumann et al. 2018; Zeevi et al. 2019; Allen et al. 2021; Langner et al. 2021). For plant studies, powerful approaches were recently proposed to associate SVs to causal genes controlling trait variation (Todesco et al. 2020; Guo et al. 2020). However, our understanding of SVs governing trait variation in microorganisms is limited by approaches focused on SNPs (Laabei et al. 2014; Pereira et al. 2020b; Singh et al. 2021). Microbial genomes are highly plastic in terms of gene content and associated SVs. GWAS based on a single reference-genome can only capture the gene content described in that single genome (Lees et al. 2016). Using a compilation of reference genomes to construct a pangenome resource that integrates a more comprehensive set of the genes present in a pathogen species shows substantial promise (Badet and Croll, 2020). The ability to integrate various types of SVs while performing association mapping will also substantially expand our understanding of microbial adaptation.

Pathogen adaptation is frequently governed by genetic determinants termed accessory genes that are not shared among all individuals of a species. Accessory genes were found to affect defense responses, virulence, drug resistance and environmental adaptation (Holt et al. 2015; Sánchez-Vallet et al. 2018; Wu et al. 2018; Zou et al. 2019). The detection of such adaptive accessory genes can be accelerated by expanding GWAS to include multiple reference genomes covering distinct segments of the gene space of a species. Additionally, single reference genome based GWAS can be confounded by gene presence/absence variation as such variation is challenging to account for (Gage et al. 2019). These shortcomings of a GWAS based on a single reference genome can be overcome by repeating the mapping across multiple reference genomes representing the pangenome of a species (Tettelin et al. 2005; Bayer et al. 2020; Gupta, 2021). Recent advances in genomics are rapidly expanding the number of microbial pathogens with such pangenome resources (Baddam et al. 2014; Liu et al. 2014; Badet et al. 2020). These resources can facilitate the identification of pathogen virulence factors as well as previously unknown anti-microbial resistance factors emerging after the application of newly designed chemical control agents (Golicz et al. 2020; Allen et al. 2021). In particular, SVs in highly repetitive regions are unlikely to be captured. This can be overcome by adopting an alignment-free approach where short reads are screened for subsequences of specific length, *i.e.* K-mers (Sheppard et al. 2013; Weinert et al. 2015). A major advantage of K-mer based analyses is the ability to capture genetic variation without depending on a reference genome, avoid SNP calling ascertainment biases or allow identifying sequence segments absent from a reference genome (Lees et al. 2016; Jaillard et al. 2018). Capturing complex SVs is particularly relevant because significant genetic variation, sometimes referred to as the “missing heritability” problem, can go undetected using traditional reference-based GWAS (Zuk et al. 2012; Rahman et al. 2018). Though their potential advantages are clear, reference-free methods to capture adaptive genetic variation remain largely unexplored in pathogenic microorganisms.

The fungal pathogen *Zymoseptoria tritici* causes septoria tritici blotch (STB), a disease that has a significant impact on global wheat production (Fones and Gurr, 2015; Torriani et al. 2015). *Z. tritici* has a highly plastic genome with 13 core chromosomes and 8 accessory chromosomes that exhibit presence-absence variation among isolates (Goodwin et al. 2011). Large effective population sizes, high gene flow and high recombination rates facilitate rapid evolution of resistance toward fungicides and virulence on resistant hosts (Croll et al. 2015; Hartmann et al. 2018, 2021; Singh et al. 2020). The pathogen population harbors substantial variation for many life history traits including growth rates, stress tolerance, melanization and reproduction on the wheat host (Dutta et al. 2021). Structural rearrangements and deletion events were found to be associated with host adaptation (Hartmann et al. 2017; Meile et al. 2018). GWAS based on single reference genomes was successful in discerning the genetic underpinnings of pathogen virulence and fungicide resistance (Hartmann et al. 2021; Singh et al. 2021). The recent pangenome constructed for *Z. tritici* based on 19 different isolates from six continents showed that the pathogen harbors a substantially larger gene repertoire than the canonical reference genome (Badet et al. 2020). Accessory genes within the species encode diverse but largely unknown functions and were likely missed in previous analyses that relied on a single reference genome. Thus, expanding GWAS beyond one reference genome will likely capture a larger fraction of genes underlying recent adaptation.

Here, we assess the performance of both reference-free and multi-reference GWAS by conducting a comprehensive mapping analysis based on a global set of *Z. tritici* populations. We screened for sources of genetic variation affecting 49 biotic and abiotic traits. Both GWAS conducted on SNP datasets mapped to 19 different reference genomes and k-mer based GWAS revealed a large number of previously missed loci contributing to trait variation. Our study provides quantitative insights how improved GWAS approaches can identify genetic variants underpinning adaptation in rapidly evolving microbial pathogens.

## Results

### A generalizable framework for conducting microbial GWAS

We performed comprehensive association mapping analyses to detect genetic variants of varying complexity underlying pathogen adaptation to different hosts and environments (**Figure 1**). We analyzed genetically diverse pathogen populations spanning the global distribution of wheat and recapitulating host diversity and climatic gradients. Isolates were phenotyped under greenhouse and laboratory conditions to assess both pathogenicity-related traits (e.g., degree of host damage, amount of spore production) and responses to abiotic stresses (e.g., fungicide, low temperature) (Dutta et al. 2021). Genetic variation in the mapping panel was assessed in two complementary ways. (1) Whole-genome sequence datasets were used to generate SNP calls on multiple reference genomes. A total of 19 telomere-to-telomere reference genomes have been assembled to capture the global diversity in structural variation (Badet et al. 2020). (2) Short reads were also used to generate 25-bp K-mer profiles for each isolate. These presence/absence K-mer tables applied to mapping populations are highly effective in capturing structural variation independent of a reference genome (Voichek and Weigel, 2020).

**Figure 1.**
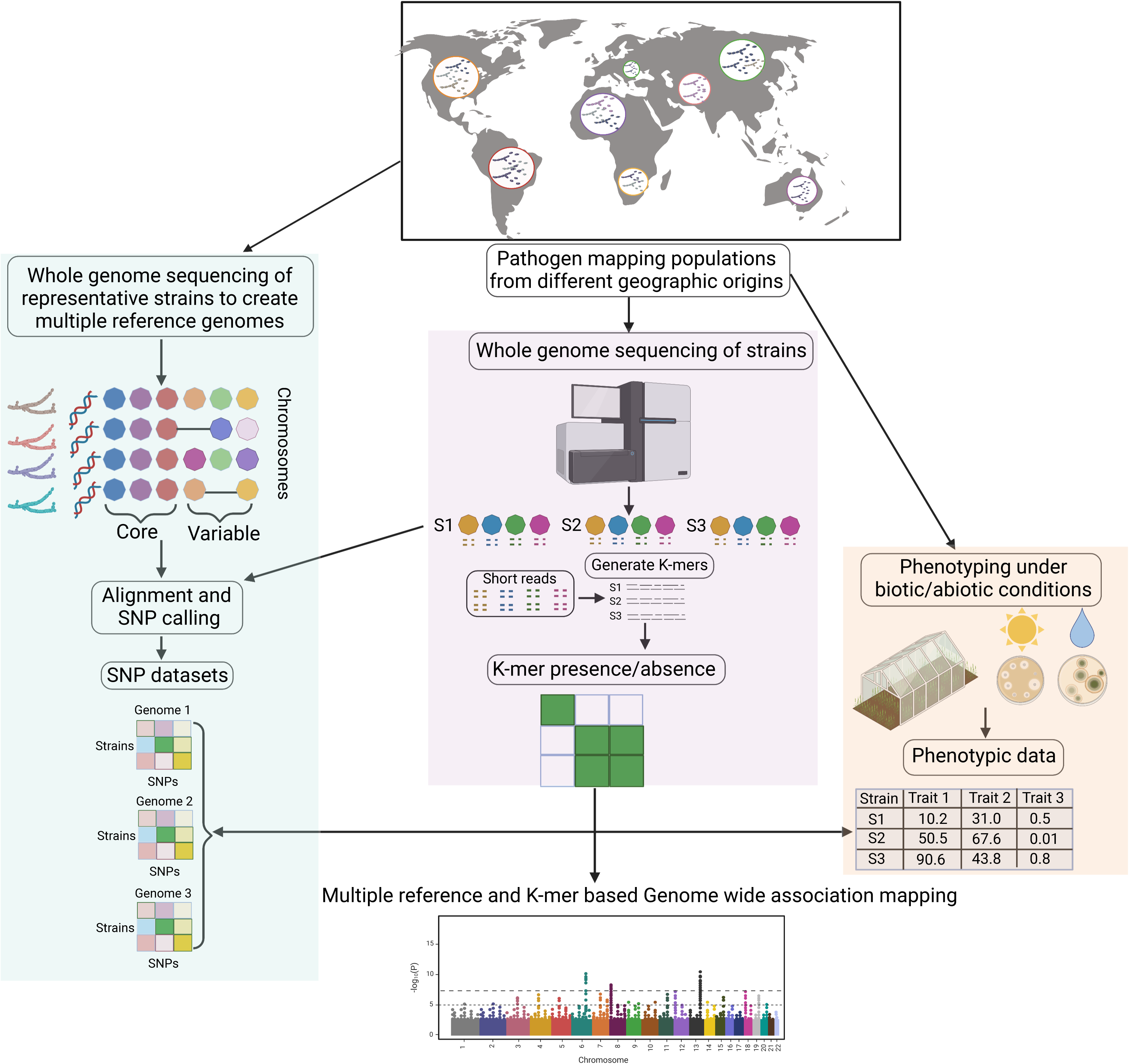
A comprehensive workflow for conducting microbial genome wide association studies (GWAS) using multiple reference genomes and K-mer data from mapping populations. Genetically diverse pathogen populations from different geographic locations are sampled to construct an association panel followed by greenhouse and laboratory phenotyping to assess heritable trait variation (right panel; Dutta et al. 2021). Chromosome-level genome assemblies of representative isolates is performed to generate reference genomes and establish a species pangenome (left panel; Badet el al. 2020). Whole genome sequencing of the association panel enables single nucleotide polymorphism (SNP) calling against multiple reference genomes and creation of K-mer presence/absence tables (middle panel). GWAS can be performed simultaneously to take advantage of SNP datasets or K-mer presence/absence tables.

### Multiple reference genome based GWAS

We performed association mapping for a total of 49 traits including measures of virulence and reproduction on twelve wheat cultivars and growth and melanization under various stress conditions such as different temperature regimes and fungicide exposure. The mapping was performed independently for SNP panels generated from each of the 19 reference genomes based on mixed linear models. We estimated the genomic inflation factor (GIF; λ), which ranged from 0.91 to 1.09 without principal components as a random effect controlling for population substructure, and from 0.70 to 1.36 when including principal components (**Supplementary Figure S1**). The multiple reference-based GWAS detected a range of significant marker-trait associations above the Bonferroni threshold (*α* = 0.05) for a total of 20 traits related to virulence, reproduction, growth rate, fungicide resistance and melanization (**Figure 2A**). We found high variability in the number of significant SNPs for the same trait depending on the choice of the reference genome SNP panel (**Figure 2A, Supplementary Table S5**). The number of significant SNPs ranged from 1-55 for pathogen virulence and reproduction on different wheat hosts depending on the reference genome. The highest number of significant SNPs were identified for virulence on landrace 1204 with the alternative reference genome KE94 (**Figure 2B**). This trait also showed the highest variance in the number of significant associations among the 19 reference genomes (**Supplementary Table S5**). The number of significant SNPs for environmental stresses ranged from 1-180 with the azole resistance trait showing the largest and most variable number of SNPs among the 19 reference genomes. The most significant SNPs for each trait explained 3-15% of the phenotypic trait variation (**Supplementary Table S6**). This suggests that numerous genes affect most trait variation in most environments, consistent with polygenic architectures for most of these traits.

**Figure 2.**
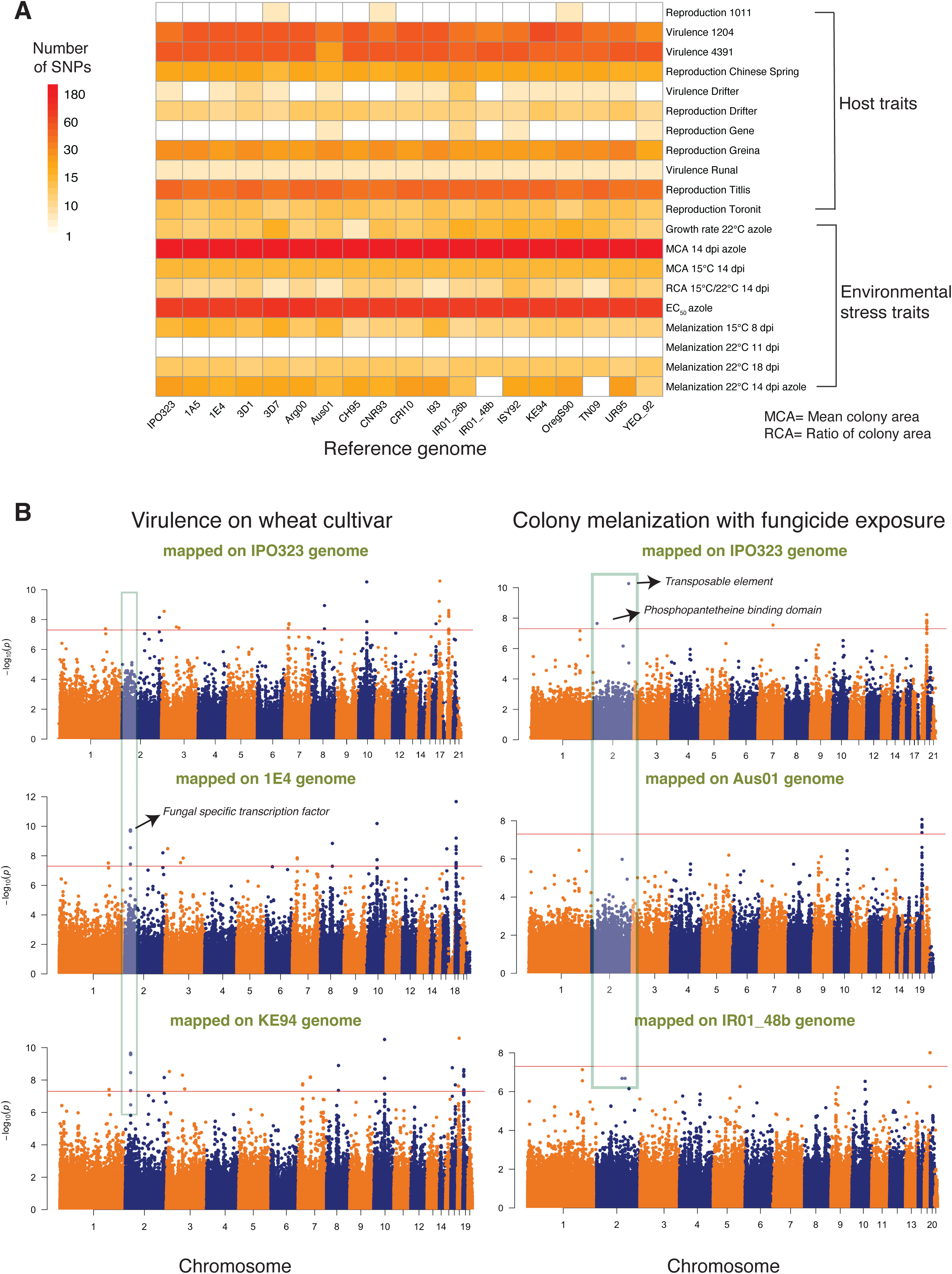
Genome wide association mapping based on 19 reference genomes for 49 pathogen traits measured under different host and abiotic conditions in *Zymoseptoria tritici*. (A) Heatmap showing differences in the number of significantly associated SNPs for each trait obtained for each reference genome. Pathogen virulence (percentage of the leaf surface covered by necrotic lesions) and reproduction (pycnidia density within lesions) were measured on 12 genetically diverse wheat lines. (B) Manhattan plots showing SNP *p*-values for two traits (pathogen virulence in the left panel and melanization in presence of fungicide in the right panel) on multiple reference genomes. The shaded gray boxes highlight differences in significant associations found when using different reference genomes. The red line indicates the Bonferroni threshold at a 5% significance level. Pathogen virulence and reproduction were measured on 12 genetically diverse wheat lines.

A substantial fraction of all significant associations could not be mapped with the canonical reference genome IPO323 (**Figure 2B**). Also, significant associations for several traits mapped in to the canonical reference genome were not found using alternative reference genomes (**Figure 2B**). This shows that multiple reference genome SNP panels can overcome limitations due to presence-absence variation and challenges in SNP calling. To analyze putative gene functions contributing to phenotypic trait variation, we extracted all the genes in close physical proximity to each SNP (< 1 kb). *Z. tritici* populations show rapid decay in linkage disequilibrium within this distance and the average distance between genes is ∼1 kb (Goodwin et al. 2011; Hartmann et al. 2017). We identified a variable number of associated genes depending on the reference genome SNP panel. The number of associated genes ranged from 54 when mapping was performed on the reference genome Aus01 to 79 on IPO323 for pathogen virulence and reproduction on different wheat hosts. The number of genes ranged from 88 (reference genome TN09) to 102 (reference genome CRI10) for environmental stress traits (*i.e.* fungicide resistance, growth rate and melanization; **Supplementary Table S7**). Based on the annotation of the canonical reference genome IPO323, the identified genes encoded a broad range of functions including major facilitator superfamily (MFS) transporters, fungal-specific transcription factors, zinc finger and protein kinase domains (**Supplementary Table S7**). Such gene functions may have specific metabolic and regulatory functions underlying pathogen adaptation (Shelest, 2008; Krishnan et al. 2018; Pereira et al. 2020b). Importantly, we detected significant SNPs near three genes encoding predicted virulence factors (*i.e.* effectors) on chromosomes 2, 5, and 7 associated with reproduction on the wheat cultivars Greina, Titlis and Chinese Spring, respectively (**Supplementary Table S7**). We also detected numerous significant SNPs for azole resistance tagging the *CYP51* gene that is known to underlie resistance to azole fungicides (Cools and Fraaije, 2012).

A challenge associated with performing multiple reference genome GWAS is to identify redundant associations across SNP panels. To estimate the extent of novel gene functions discovered through the expansion of the reference genome SNP panels, we performed a saturation analysis based on orthology information. For each gene with a significant association, we assessed whether any ortholog identified in a different reference genome was already tagged (*i.e.* is a member of the same orthogroup). We randomly selected subsets of the reference genome SNP panels and counted the number of unique orthogroups with significant associations for groups of traits. We observed a near-linear increase in the number of unique orthogroups with significant associations with an increasing number of reference genome panels (**Figure 3**). The most substantial increase was observed by including a second reference genome panel. Beyond two reference genome panels, the benefits for each additional reference genome SNP panel decreased slightly. This shows that a substantial fraction of the genetic factors contributing to adaptation to host, and environmental stress factors cannot be identified from a single reference genome SNP panel. Fungicide resistance related traits show the highest number and fastest gain in significantly associated orthogroups with additional reference genome SNP panels. Pathogen virulence and reproduction showed intermediate increases in significantly associated orthogroups and melanization showed the slowest increase in significantly associated orthogroups. Overall, including multiple reference genome SNP panels substantially expands the spectrum of identifiable genetic factors (**Supplementary Figure S3**).

**Figure 3.**
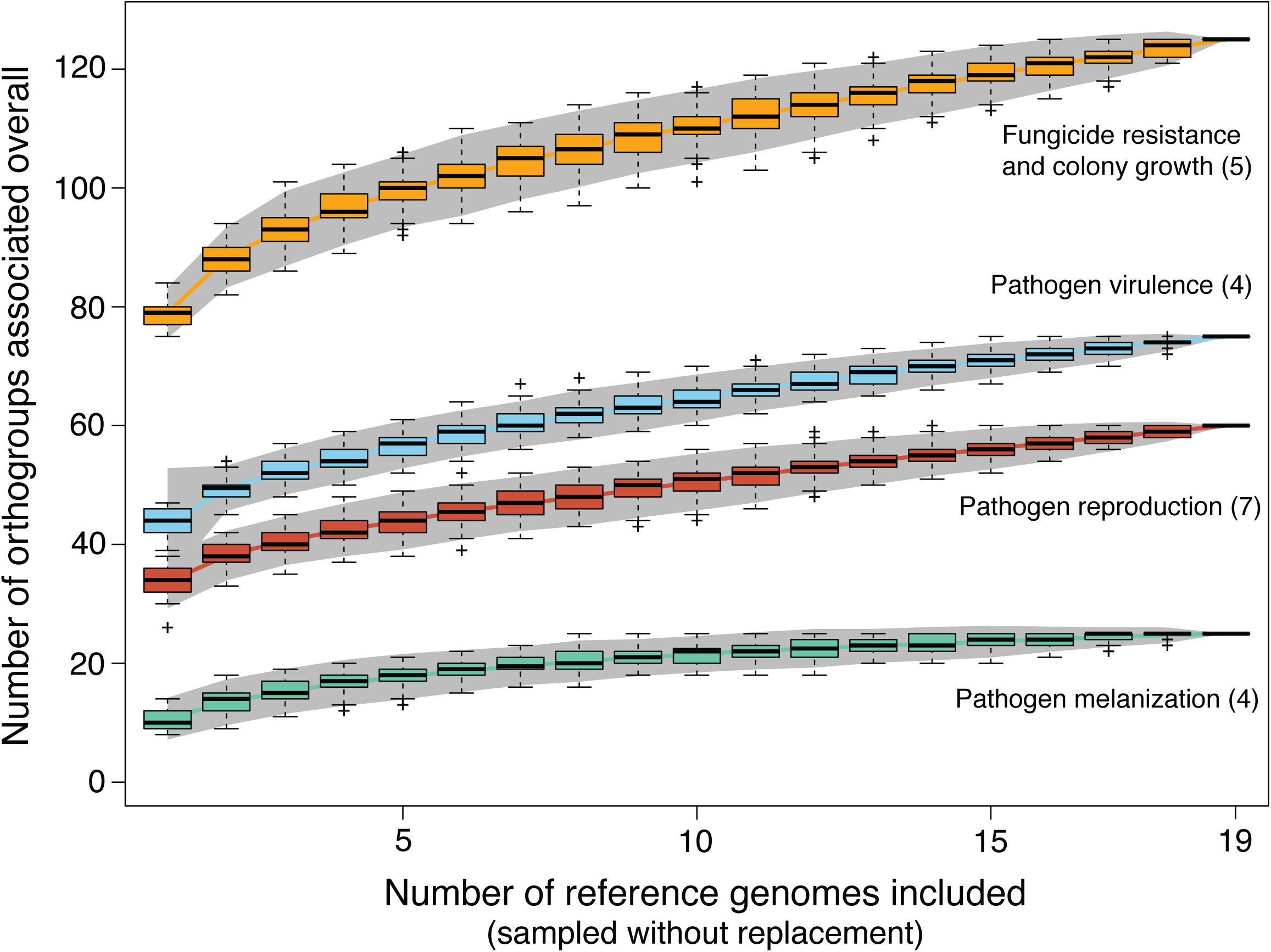
Accumulation curves for the total number of distinct genes (identified by orthogroups within the species) associated with GWAS for different traits as a function of the number of reference genomes analyzed. Mapping outcomes are shown for different groups of traits. The numbers in parentheses indicate the number of traits included in each category. Pathogen virulence (percentage of the leaf surface covered by necrotic lesions) and reproduction (pycnidia density within lesions) were measured on 12 genetically diverse wheat lines.

### K-mer approach to uncover additional sources of genetic variation

To further expand our survey of structural variation potentially associated with trait variation, we performed reference-free GWAS on the same trait dataset using 25-bp K-mers generated from whole genome sequencing data. We identified a total of ∼55 million K-mers of which 7,111,640 were detected in at least five isolates. We estimated K-mer based heritability to contrast with SNP-based heritability from Dutta et al. (2021). For pathogen virulence, K-mers explained a higher proportion of phenotypic variance compared to the SNP-based estimates (**Figure 4A**). A similar trend of increased heritability accounted by K-mers was observed for all other traits as well (**Supplementary Figure S2A, 2B, 2C**). The heritability for virulence ranged from 0 to 0.84 (standard error, SE=0.08) with an average of 0.6 (SE=0.16) compared to 0.35 (SE=0.14) based on SNPs. Heritability for reproduction traits ranged from 0.73 (SE=0.13) to 0.96 (SE=0.01) with an average of 0.86 (SE=0.06) compared to SNP-based heritabilities with an average of 0.65 (SE=0.1). The average heritability for environmental stress factors (i.e., fungicide resistance, growth rate and melanization at different temperatures) was 0.7 (SE=0.18) compared to 0.51 based on SNPs (SE=0.18). Consistent with the high heritability estimates, the K-mer GWAS yielded numerous K-mers above the permutation-based significance threshold (*α* = 0.05) for 33 out of 49 phenotypic traits. The number of significant K-mers ranged from 3-2066 for pathogen virulence, from 3-640 for pathogen reproduction, from 3-166 for pathogen melanization, and from 9-3606 for fungicide resistance and growth-related traits.

**Figure 4.**
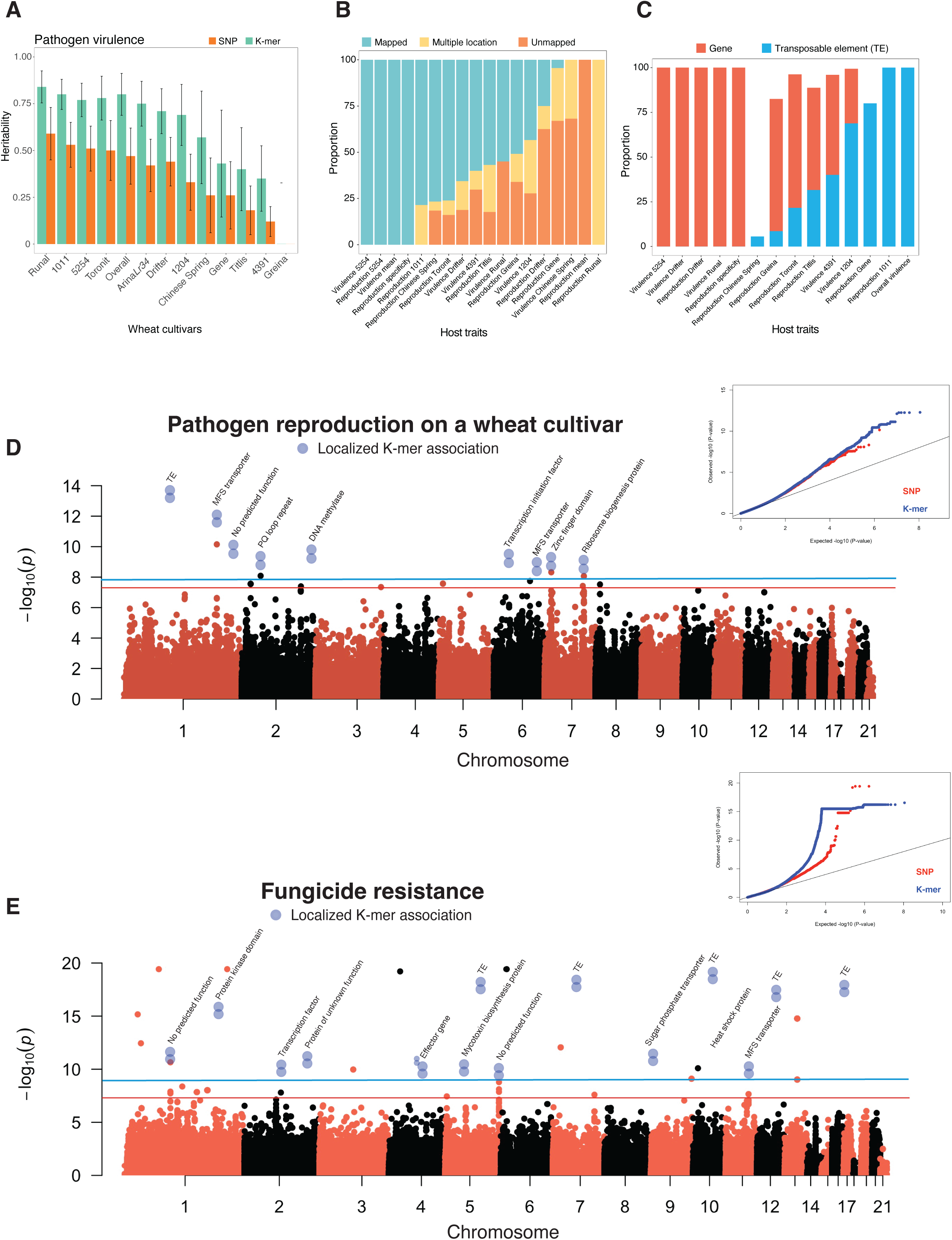
K-mer GWAS on 49 life-history traits based on a K-mer presence/absence table for all 145 *Zymoseptoria tritici* isolates. (A) Comparison of heritability estimates for pathogen virulence (percentage of the leaf surface covered by necrotic lesions) based on SNPs (for the reference genome IPO323) and K-mers. Both SNP-based and K-mer-based heritability were estimated by following a genome-based restricted maximum likelihood (GREML) approach. Standard errors are indicated by error bars (B) Alignment of significantly associated K-mers against the reference genome (IPO323) show the proportion of K-mers having a unique mapping position, multiple locations, or no unambiguous mapping position in host-related traits *i.e.* pathogen virulence and reproduction (pycnidia density within lesions). (C) Proportion of significant K-mers with a unique mapping position in the reference genome either tagging a gene or a transposable element for host-related traits. (D, E) Manhattan plots showing significant K-mer associations with pathogen reproduction and fungicide resistance together with quantile-quantile plots for *p*-value comparisons. Manhattan plots were created from SNP-based GWAS and blue dots represents the significant K-mer associations with the K-mers being uniquely mapped to a location in the reference genome. The two blue dots represent individual K-mers with significant associations. The red and blue lines indicate the Bonferroni and permutation-based significance threshold at 5% level for SNPs and K-mers, respectively. Pathogen virulence and reproduction were measured on 12 genetically diverse wheat lines. Overall virulence and reproduction represent the average value of the respective trait measured on 12 genetically diverse wheat lines. Reproduction specificity was estimated based on the adjusted coefficient of variation of mean reproduction across 12 genetically diverse wheat lines. Higher specificity suggests affinity to certain hosts for maximizing reproductive fitness.

To identify gene functions mapped through K-mer GWAS, we searched K-mer sequences in the canonical reference genome IPO323 (**Figure 4B, Supplementary Figure S2D**). We found a substantial fraction of significant K-mers tagging either a transposable element (TE) or a gene in the *Z. tritici* genome (**Figure 4C, Supplementary Figure S2E**). For host-related traits (**Figure 4B**), an average of 63.6% of all significant K-mers tagged a gene compared to 32.1% tagging a TE. In contrast, the proportions of significant K-mers tagging a TE or a gene were roughly inverted (59.17% *vs.* 34.6%) for environmental stress traits (**Supplementary Figure S2D)**. Furthermore, for the majority of the traits, the K-mer with the highest *p*-value tagged a TE (**Figure 4D, 4E**). The high proportion of K-mers mapping to a TE suggests that active transposition has contributed significantly to phenotypic variation in *Z. tritici*. Additionally, the K-mer GWAS discovered a large number of not previously identified genes associated with both host-related and environmental stress traits (**Figure 4D, 4E; Supplementary Figure S3**). The K-mer tagged genes encoded a broad range of functions including a transcription factor, MFS transporters, and peptidases as well as effector candidates (**Figure 4D, 4E, Supplementary Table S8)**.

We analyzed in detail how the K-mer approach expanded the discovery of loci compared to SNP-based GWAS. We focused on the key azole resistance gene *CYP51* (**Figure 5A**). We found 294 K-mers above the 5% significance threshold on chromosome 7 associated with *CYP51* (*Zt09_07_00450*) for the resistance trait EAM_14_dpi_azole. All the K-mers could be located to a unique position on the chromosome. The K-mer *p*-values tagging this gene were lower than the SNP *p*-values (**Figure 5B**). Nearly all (293/294) K-mers were located in the upstream region of the gene spanning between the positions 1,446,325 and 1,446,893 bp. The K-mer presence/absence among isolates were in full linkage disequilibrium (**Figure 5C, 5D**; *r^2^*=1). One additional K-mer localized (1,447,308 bp) to the fourth and largest exon of the gene and showed lower linkage disequilibrium (*r^2^*=0.48) with the other K-mers. Most of the isolates from Switzerland (71.1%) and a few from Israel (10%) carried the K-mers associated with increased azole resistance (**Figure 5E; 5F**). We expanded our analyses of K-mer associations to virulence traits (**Figure 6A**). We discovered 11 significant K-mers on chromosome 7 (from 1,897,941-1,897,951 bp) for virulence on the cultivar Runal. The tagged gene was previously identified through QTL mapping and encodes a virulence factor termed *Avr3D1.* No SNPs in the same region passed the Bonferroni significance threshold (**Figure 6B**). All K-mers were located in the largest exon and all but one was in full linkage disequilibrium with each other (**Figure 6C, 6D**). The K-mer with lower linkage disequilibrium to the other K-mers was primarily detected in isolates of the Israel population (**Figure 6E**). The isolates carrying the significant K-mers produced less leaf damage (**Figure 6F**).

**Figure 5.**
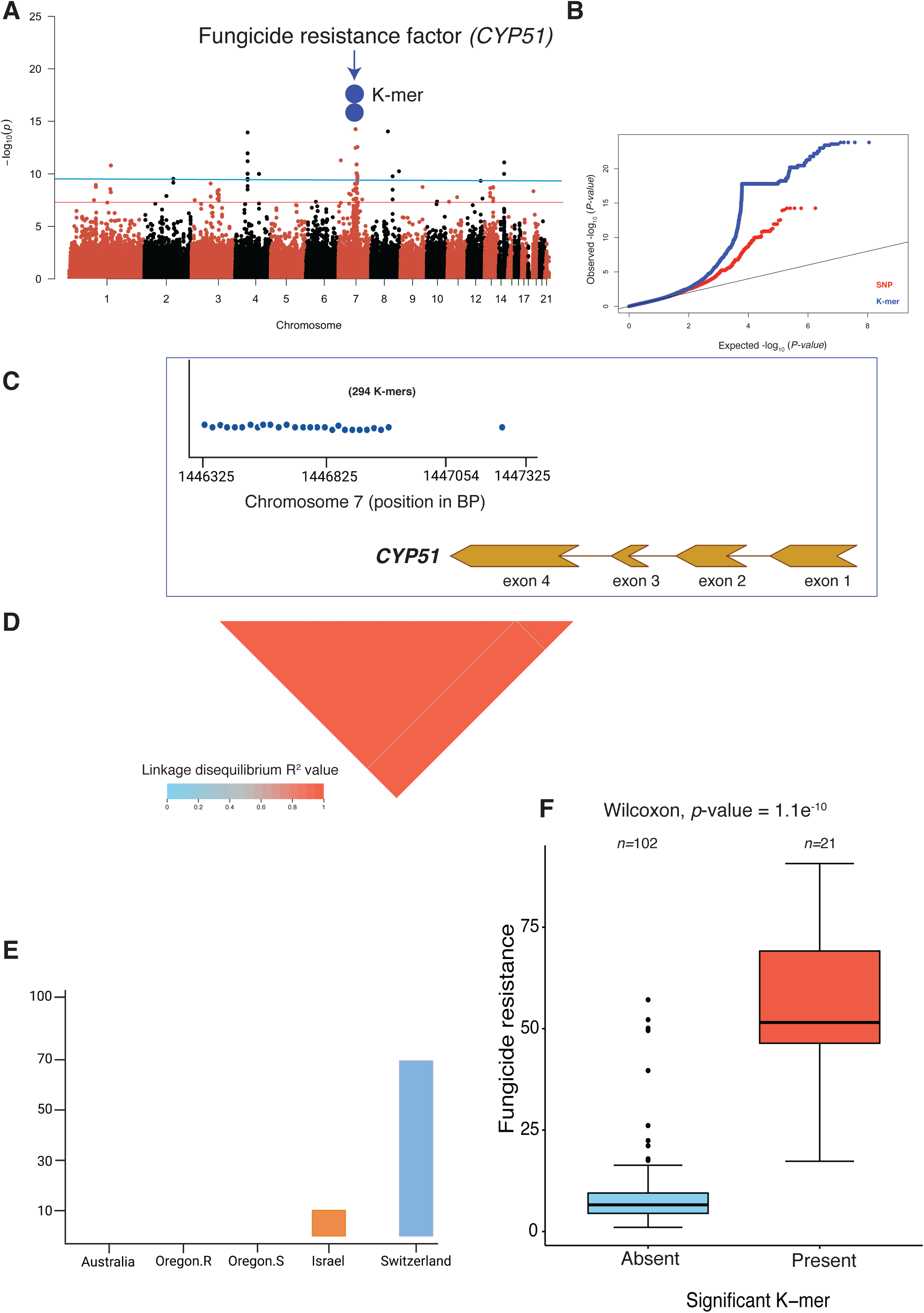
Analysis of K-mer GWAS identifying causal genes underlying major phenotypes in *Zymoseptoria tritici*. (A) Manhattan plot showing significant K-mers associated with fungicide resistance. The two blue dots represent all 294 significant K-mers with a unique genomic position on chromosome seven tagging the *CYP51* gene encoding the target of azole fungicides. The red and blue lines show the Bonferroni and permutation-based significance threshold (*α*=0.05) for SNP and K-mer GWAS, respectively. (B) Quantile-Quantile plot showing the *p*-value comparison between SNPs and K-mer based GWAS. (C) Physical position of 294 significant K-mers mapped to unique positions on chromosome seven associated with the fungicide resistance gene *CYP51*. (D) Linkage disequilibrium (LD) heatmap showing the pairwise *r^2^* value among 294 significant K-mer presence/absence genotypes associated with the *CYP51* gene. (E) Proportion of isolates from different populations carrying significant K-mers that tagged *CYP51*. (F) Boxplot showing fungicide resistance levels in isolates with presence of the K-mers associated with the *CYP51* gene.

**Figure 6.**
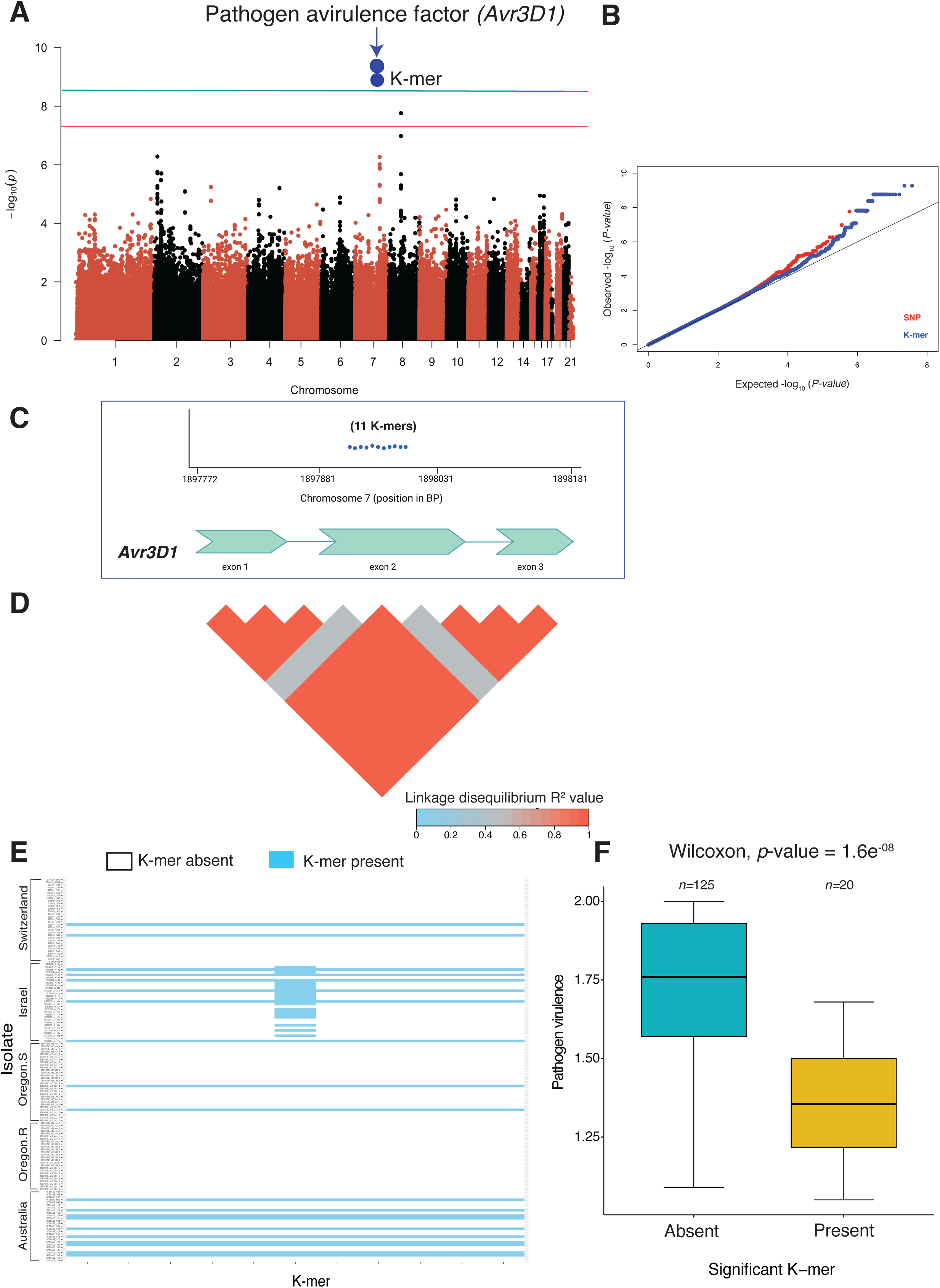
K-mer based GWAS recovered a known effector gene in *Zymoseptoria tritici* with a higher statistical power than SNP-based GWAS. (A) Manhattan plot showing significant K-mers associated with pathogen virulence on the wheat cultivar Runal. The two blue dots represent all 11 K-mers uniquely mapping to positions on chromosome seven and tagging the avirulence gene *Avr3D1* encoding an effector protein. The red and blue lines indicate the Bonferroni and permutation-based significance threshold (*α*=0.05) for SNP and K-mer GWAS, respectively. (B) Quantile-Quantile plot showing the *p*-value comparison between SNPs and K-mers. (C) Physical position of 11 uniquely mapped K-mers on chromosome seven associated with *Avr3D1*. (D) Linkage disequilibrium (LD) heatmap showing the pairwise *r^2^* value among 11 significant K-mers associated with *Avr3D1*. (E) Presence/absence pattern of 11 significant K-mers associated with *Avr3D1* in five *Z. tritici* populations. The continuous horizontal blue line indicates isolates containing all the significant K-mers. (F) Boxplot showing pathogen virulence (percentage of the leaf surface covered by necrotic lesions) on the wheat cultivar Runal in isolates with or without the significant K-mers associated with *Avr3D1*.

## Discussion

Here, we report the most comprehensive assessment of association mapping performance to date for microbial pathogens to unravel genetic determinants of phenotypic trait variation. We find that expanding association mapping to include multiple reference genome SNP datasets provides a near linear increase in the number of additional loci detected by GWAS. Performing a reference-free GWAS approach using K-mers similarly boosted the power to uncover genetic variation underlying important traits. The extensive gains in the power of GWAS analyses that take into account structural variation reveals a greater proportion of the complexity inherent in adaptive genetic variation within microbial species.

SNP-based GWAS based on a single reference genome dataset have been successful in describing the genetic basis of complex pathogen traits (Mohd-Assaad et al. 2016; Pereira et al. 2020b; Caseys et al. 2021; Singh et al. 2021). By expanding the number of reference genome SNP datasets used for GWAS, we identified substantially more independent loci than what was previously identified using the same phenotype dataset (Dutta et al. 2021). The number of loci associated with most trait variation increased almost linearly with the addition of reference genome SNP datasets. Such an increase is striking given the fact that most traits are thought to be significantly constrained by stabilizing selection and have a conserved genetic basis (*e.g.* growth, melanization, reproduction; Steffansson et al. 2014, Qin et al. 2016, Pereira et al. 2020a). Stabilizing selection tends to reduce shared additive genetic variation between populations and closely related species, which ultimately reduces phenotypic variation (Yair and Coop, 2021). Pathogen trait expression is expected to stabilize at an optimal level due to genetic trade-offs (Bonneaud et al. 2020; Dutta et al. 2021). Climatic conditions and host genotype turnover may lead to rapid shifts in selection pressures. Hence, there should also be turnover in the genes underlying adaptation to previous environmental conditions. The *Z. tritici* pathogen model may be an outlier given the maintenance of very large population sizes, high gene flow and extensive chromosomal polymorphism (Hartmann et al. 2018; Badet and Croll, 2020). The near-linear increase in associated loci may also be explained, at least in part, by the use of a highly diverse, global panel of reference genomes. The reference genome isolates originating from six continents stem from populations that likely experienced divergent selection pressure from locally adapted hosts and local climatic conditions. Overall, we show that including a broad set of reference genome SNP datasets efficiently overcomes limitations imposed by using a single reference genome. Such limitations often stem from ascertainment bias in SNP calling and genetic distance between the reference genome and mapping populations (Valiente-Mullor et al. 2021). A particular concern is that a single reference may not represent the full catalog of gene functions relevant for adaptation in the species pool (Golicz et al. 2020). For instance, missed associations for genes that are absent from a reference genome may underpin an adaptive advantage in a specific ecological context and/or geographic region (Lassalle et al. 2015; Gori et al. 2020).

We find that accounting for genetic variation using K-mers instead of SNPs explains more genetic variation (*i.e.* gives a higher heritability). This implies that significant phenotypic variation is explained by genetic factors located in genomic regions that are difficult to access using SNPs. Such genetic variants are likely to be found in non-coding and TE-rich regions. Such variants may be in accessory genomic regions absent from the reference genome and not easily assessed through SNP calling. Missing heritability in human traits has been recovered by including rare genetic variants (Wainschtein et al. 2021). We show that incorporating genetic variants other than SNPs in plant pathogen GWAS increases trait heritability as well. We also found K-mers in extremely polymorphic regions of the core genome such as the regions surrounding the genes *CYP51* and *Avr3D1*. Recent studies have shown that SVs such as chromosomal rearrangements and copy number variations contribute to adaptive evolution in pathogens (Peter et al. 2018; Firrao et al. 2018; Badet et al. 2021). The K-mer approach broadly revealed three classes of loci: (1) loci previously identified by SNP-based GWAS, (2) gene functions that were not identified through SNP-based GWAS but have independent evidence for their contribution to phenotypic trait variation (*i.e. CYP51* and *Avr3D1*) and (3) previously unknown gene functions including genes encoding effector candidates for host manipulation and genes encoding detoxification functions (*e.g.* MFS transporters). The K-mer approach for GWAS has been successfully implemented for plants (Voichek and Weigel, 2020) and bacteria (Lees et al. 2016; Young et al. 2019). Here we provide strong evidence that such reference-free GWAS can also be successfully performed in eukaryotic microbial pathogens.

Genetic variation in plant pathogens is characterized by high degrees of functionally relevant polymorphism as well as genomic plasticity underpinning accessory genes (Ehrlich et al. 2005; Hammond et al. 2020; Badet and Croll, 2020). Beyond this, we found substantial complexity in the genes underlying the expression of the same trait under different environmental conditions. Working with such highly diverse pathogen populations poses serious challenges for selecting appropriate reference genome resources. Here we show that GWAS conducted on multiple reference genome SNP datasets and using reference-free approaches effectively compensates for this genetic diversity. This is supported by our recovery of known causal loci for specific phenotypes, including loci missed by previous GWAS, as well as a general improvement in heritability for all traits. Further refinements of our approach should integrate recent developments such as pangenome graphs that might alleviate limitations of studies based on SNPs and single reference genomes. Leveraging a multitude of GWAS signals following our combinatorial approach is likely to significantly advance our mechanistic understanding of pathogen emergence and adaptation.

## Methods

### Fungal material

A collection of 145 *Z. tritici* isolates sampled independently from four different wheat fields was used in this study. The field isolates were sampled between 1990 and 2001 from four different countries (Zhan et al. 2005): Australia (*n*=27), Israel (*n*=30), Switzerland (*n*=32) and USA (Oregon.R, *n*=26; Oregon.S, *n*=30). The two Oregon populations were sampled from the wheat cultivar Madsen (moderately resistant) and Stephens (susceptible), growing simultaneously in the same field. Clones were removed from the field populations so that the analyzed panel comprises only strains with unique genotypes. Blastospores of each isolate were preserved in either 50% glycerol or anhydrous silica at −80°C.

### Phenotyping for host infection traits

Datasets on virulence and reproduction for each pathogen strain were previously established by Dutta et al. (2020) (**Supplementary Table S1**). Virulence and reproduction were measured on 12 genetically different wheat cultivars displaying varying degrees of resistance and susceptibility to STB. The wheat panel included six commercial varieties (Drifter, Gene, Greina, Runal, Titlis, Toronit), a back-cross line (Arina*Lr34*) and five landraces (1011, 1204, 4391, 5254, Chinese Spring). Four of the landraces (1011, 1204, 4391, 5254) came from the Swiss National Gene Bank (www.bdn.ch). Detailed phenotyping protocols are described in Dutta et al. (2020). Briefly, three seeds of each cultivar were planted in a six-pot strip arrayed in a 2 × 3 pattern. Due to space limitations, the experiment was conducted in two stages, each including six cultivars. All plants were maintained in a greenhouse chamber at 22 °C (day) and 18 °C (night) with 70% relative humidity (RH) and a 16-h photoperiod. Blastospores of each isolate were inoculated using an airbrush spray gun until run-off on two-week-old seedlings to initiate the infection process. In both stages, the inoculations were repeated separately three times to generate three biological replications in separate greenhouse chambers. All inoculated second leaves were collected between 19-26 days post inoculation (dpi) and fixed on QR-coded A4 paper for scanning. The scanned images were analyzed using automated image analysis (AIA; Karisto et al. 2018) to generate quantitative data on the amount of damaged leaf tissue (*i.e.* virulence) and the density of pathogen fruiting bodies called pycnidia produced within the damage area (*i.e.* reproduction).

### Phenotyping for growth and stress-related traits

In vitro traits comprised fungal growth rate (mm per day), thermal sensitivity, mean colony area, fungicide resistance and melanization measured at different temperatures with or without fungicide (**Supplementary Table S2)** following previously described phenotyping protocols (Lendenmann et al. 2014, 2015, 2016; Mohd-Assaad et al. 2016). Briefly, after revival from long-term storage, each isolate was cultured on Petri dishes filled with yeast malt sucrose agar (4 g/L yeast extract, 4 g/L malt extract, 4 g/L sucrose, 50 mg/L kanamycin) for 4-5 days at 18 °C. Blastospore solutions were diluted using sterile water to a final concentration of 200 spores/ml using KOVA counting slides (Hycor Biomedical, Inc., Garden Grove, CA, USA). Petri dishes containing potato dextrose agar (PDA, 4 g/L potato starch, 20 g/L dextrose, 15 g/L agar) were inoculated with 500 µl of the spore solution. Inoculated plates were maintained at 15 °C (cold treatment) or 22 °C (control treatment) at 70% RH. Images were captured with a digital camera at 8, 11 and 14 days post inoculation (dpi) to generate five technical replicates. The photographs were analyzed using AIA macros in ImageJ as described in Lendenmann et al. (2014) to measure colony growth. The estimates of colony growth rate for each isolate were obtained by fitting a general linear model over three time points by taking the mean colony radii from 45 colonies. The growth rate ratio between colonies growing at 15 °C or 22 °C, or on 22 °C PDA plates with or without propiconazole (Syngenta, Basel, Switzerland; 0.05 ppm) were expressed as temperature and fungicide sensitivity at 14 dpi, respectively. Fungicide resistance was also quantified on microtiter plates by growing 100 µl spore solutions at a concentration of 2.5 × 10^4^ spores/ml of each isolate on 100 µl Sabouraud-dextrose liquid media (SDLM; 20 g/L dextrose, 5 g/L pancreatic digest of casein, 5 g/L peptic digest of animal tissue; Oxoid, Basingstoke, UK) with 12 different concentrations of propiconazole (0, 0.00006, 0.00017, 0.0051, 0.0086, 0.015, 0.025, 0.042, 0.072, 0.20, 0.55, 1.5 ppm propiconazole). Plates containing fungal spores amended with the fungicide of each isolate were gently shaken for one minute, sealed and incubated in the dark for four days at 22 °C with 80% RH. Three technical replicates of each isolate were performed. Fungal growth was estimated with an ELISA plate reader (MR5000, Dynatech) by examining the optical density (OD) at 605 nm wavelength. We estimated the EC_50_ value (concentration at which the growth was reduced by 50%) for each isolate using dose-response curves across the varying fungicide concentrations using the drc v.3.0-1 package (Ritz et al. 2015) in the R-studio (R Core Team, 2014). Melanization of each isolate was measured at 8, 11, 14 and 18 dpi during growth at 15°C, 22°C and at 22°C with 0.05 ppm propiconazole. We measured the mean gray value of fungal colonies from replicated plates for each isolate ranging from 0 (black) to 255 (white) for each time point. To provide a more intuitive interpretation of melanization, each mean gray value was subtracted from 255 to transform the original melanization scale to range from 0 (white) to 255 (black).

### Read mapping and single nucleotide polymorphism calling

We used publicly available raw Illumina whole genome sequences of 145 *Z. tritici* isolates (**Supplementary Table S3**; Dutta et al. 2021). Trimmomatic v.0.36 (Bolger et al. 2014) was used with the following settings (illuminaclip = TruSeq3-PE.fa:2:30:10, leading = 10, trailing = 10, slidingwindow = 5:10, minlen = 50) to trim off low-quality reads and remove adapter contamination from each isolate. Trimmed sequence data from all isolates were aligned to the *Z. tritici* reference genome IPO323 (Goodwin et al. 2011) using Bowtie2 v.2.3.3 with the option “-- very-sensitive-local” (Langmead et al. 2009). We removed PCR duplicates from the alignment (.bam) files by using the MarkDuplicates module in Picard tools v.1.118 (http://broadinstitute.github.io/picard). Single nucleotide polymorphism (SNP) calling and variant filtration steps were performed using the Genome Analysis Toolkit (GATK) v.4.0.1.2 (McKenna et al. 2010). We performed SNP calling for all 145 *Z. tritici* isolates independently using the GATK HaplotypeCaller with the command “-emitRefConfidence GVCF; -sample_ploidy 1” (*Z. tritici* is haploid). Then, GenotypeGVCFs was used to conduct joint variant calls on a merged gvcf variant file with the command -maxAltAlleles 2. SNPs found only in the joint variant call file were retained. As recommended by GATK Best Practices, we performed hard filtering of SNPs based on quality cut-offs using the GATK VariantFiltration and SelectVariants tools. Variants matching any of the following criteria were removed: QUAL < 250 (overall quality filter); QD < 20.0 (avoiding quality inflation in high-coverage regions); MQ < 30.0 (avoid calls from ambiguously mapped reads); −2 > BaseQRankSum > 2; −2 > MQRankSum > 2; −2 > ReadPosRankSum > 2; FS > 0.1. Using this procedure, the genotyping accuracy was shown to be high and congruent with an alternative SNP caller (Hartmann et al. 2018). We retained a genotypic call rate of ≥80% and minor allele frequency (MAF) > 5% to generate a final SNP dataset containing 883,207 biallelic SNPs based on the reference genome IPO323. We repeated the SNP calling and filtering procedure separately for 18 additional fully assembled *Z. tritici* genomes from Badet et al. (2020). The number of biallelic SNPs called on the 18 additional reference genomes ranged from 827,851 (genome TN09) to 883,119 (genome I93; **Supplementary Table S4**).

### SNP-based genome-wide association mapping

Log-transformed least-square means for each isolate × environment combination including 49 traits were obtained from Dutta et al. (2021) to conduct genome-wide association (GWAS) mapping. We used a mixed linear model (MLM) approach implemented in the program GEMMA v.0.98 (Zhou and Stephens, 2012) to perform GWAS on all the traits. MLMs control for genetic relatedness and population structure (Kang et al. 2008; Zhang et al. 2012). Prior to GWAS, we converted all 19 SNP datasets (one per reference genome) into PLINK “.bed” format to perform principal component analyses (PCA) using the “--pca” command in PLINK v.1.90 (Purcell et al. 2007). To account for genetic relatedness among isolates, a centered genetic relatedness matrix (GRM) for each SNP dataset was constructed using the option “-gk 1” in GEMMA by considering all genome-wide SNPs. As both PCA and GRM can efficiently control for *p*-value inflation, we estimated genomic inflation factors (GIF, λ; Devlin and Roeder, 1999) to make decisions on whether PCs should be included in the GWAS models as covariates or not. The GIF for each trait was estimated as λ = M/E, where M is the median of the observed chi-squared test statistics and E is the expected median of the chi-squared distribution (Yang et al. 2011). The distribution of all SNP effects follows a one degree of freedom chi-square distribution under the null hypothesis with a median of ∼0.455, which can be inflated by discrepancies in allele frequencies caused by population structure, genetic relatedness, and genotyping errors. The inflation is proportional to the deviation from the null hypothesis. When the fitted GWAS model efficiently accounts for such systematic deviations, the λ value is close to 1. Therefore, depending on the λ value, the reference genome based GWAS were performed using either LMM+K or LMM+K+PC, where K is the GRM used as a random effect and the first three PCs were used as fixed covariates. We used the following LMM model in GEMMA:

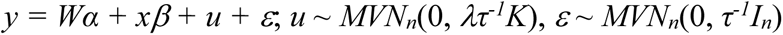

where *y* represents a vector of phenotypic values for *n* individuals; *W* is a matrix of covariates (fixed effects with a column vector of 1 and the first three PCs), *α* is a vector of the corresponding coefficients including the intercept; *x* is a vector of the genotypes of the SNP marker, *β* is the effect size of the marker; *u* is a vector of random individual effects; *ε* is a vector of random error; *τ^-1^* is the variance of the residual errors; *λ* is the ratio between the two variance components; *K* is the *n* × *n* genetic relatedness matrix and *I_n_* is an *n* × *n* identity matrix and *MVN_n_* represents the multivariate normal distribution. We set the MAF to 5% with a maximum of 50% missing values with the option “-miss 0.5”. SNP *p*-values were estimated following a likelihood ratio test in GEMMA. We used the stringent Bonferroni threshold (*α* = 0.05; *p* = *α* / total number of SNPs) to define a SNP significantly associated with a phenotype. The proportion of phenotypic variance explained by the most significant SNPs was estimated by *2f(1-f)a^2^*, where *f* is the minor allele frequency and *a* is the standardized coefficient (Gudbjartsson et al. 2008). To obtain the standardized coefficient for each SNP, we estimated the standardized regression coefficient applying a linear regression model with the “standard-beta” option implemented in PLINK v.1.9. We restricted this analysis only to the canonical reference genome IPO323. To identify genes close to significantly associated SNPs in one of the reference genomes (Badet et al. 2020), we used the BEDtools v.2.29.0 (Quinlan and Hall, 2010) *closest* command. We further investigated patterns of linkage disequilibrium (LD) in the genomic regions with the most significantly associated SNPs. All possible SNP pairs in 5 kb windows were analyzed using the “--hap-r2” command in vcftools. To visualize the *r^2^* values, heatmaps for each locus were generated using the R package LDheatmap v.0.99-7 (Shin et al. 2006). We created a heatmap summarizing the number of significant SNPs passing the Bonferroni threshold for each trait and each genome using the R package *pheatmap* (Kolde, 2012).

### K-mer based genome-wide association mapping

We performed K-mer based GWAS on all 49 traits in the panel of 145 *Z. tritici* isolates following the methodology described in Voichek and Weigel (2020). This approach uses raw sequencing reads of specific length and was designed for settings where a reference genome is lacking or to account for structural variation. K-mers of 25 bp length were counted with and without canonization, sorted and listed in a textual format for each isolate separately. K-mer canonization refers to storing K-mers and their reverse-complement for generating presence/absence patterns since these sequences are indistinguishable (Voichek and Weigel, 2020). K-mer length has an impact on the number and accuracy of K-mers. For small genomes of the size of *Z. tritici,* 25-bp K-mers are recommended (Voichek and Weigel, 2020). K-mers were filtered based on the presence/absence patterns among isolates with a 5% MAF and compressed into a presence/absence table for running GWAS. There were 55,758,186 unique K-mers generated from 145 isolates. Prior to GWAS, a GRM was estimated with EMMA (Efficient Mixed-Model Association) that comprised an identity-by-state (IBS) matrix under the assumption that each K-mer has a small, random effect on the phenotype. GWAS was performed by using an LMM+K model in GEMMA with the likelihood ratio test to estimate *p*-values. A K-mer was considered to be significant when the *p*-value passed the permutation-based threshold as described in Voichek and Weigel (2020). The pairwise LD among significant K-mers for each trait was estimated by converting the K-mer presence/absence table containing all the K-mers into PLINK format and using the command “-- r2” in PLINK. We attempted to map all the significant K-mers for each trait to the *Z. tritici* reference genome IPO323 using the short-read aligner bowtie v1.2.2 (Langmead and Salzberg, 2012) with the command “-a --best –strata”. We used the center position of the K-mer alignment to the reference genome as a coordinate to inspect nearby features using BEDtools. If no significant K-mer could be mapped to the reference genome, we retrieved the isolates carrying the specific K-mer and used the paired-end raw sequencing reads to detect the origin of the K-mer. These paired-end reads were then aligned to the canonical reference genome IPO323 using Bowtie2 v.2.3.3 (Langmead and Salzberg, 2009).

### Heritability estimation using SNPs and K-mers

We estimated SNP-based heritability on multiple reference genomes and K-mer-based heritability following the same procedure described in Dutta et al. (2021). Briefly, the phenotypic data of each trait and the GRM representing the additive effect of all genome-wide SNPs from the canonical reference genome IPO323 and K-mers were included in a genome-based restricted maximum likelihood (GREML) approach using the genome-wide complex trait analysis (GCTA) tool v.1.93.0 (Yang et al. 2011) to estimate heritability. GRMs for reference genome SNP datasets and the K-mer presence/absence table (converted into PLINK format) were estimated following a normalized identity-by-state method and fitted as a random factor in the model to estimate the proportion of phenotypic variance for each trait. The following formula from Yang et al. (2011) was used to estimate the relatedness between two individuals:

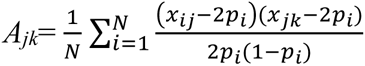

Where *x_ij_* is the number of copies of the reference allele for the *i^th^* SNP of the *j^th^* individual and *p_i_* is the frequency of the reference allele and *N* is the number of SNPs. Here, the GRMs were constructed using all genome-wide SNPs and K-mers irrespective of the nature of their relationship with the phenotype, thus indicating the approximated genetic similarities at causal loci and the accuracy of the heritability estimates.

### Pangenome analyses

We generated accumulation curves to estimate the gain in additional loci from performing GWAS on more than one reference genome. For this, we retrieved for each GWAS based on SNPs mapped to a particular reference genome the set of genes within 1 kb distance with significantly associated SNPs. Then, we matched the set of associated genes among genomes using within-species gene orthology information (Badet et al. 2020) to determine whether genes belong to the same orthogroup. We used a sampling procedure (without replacement) among reference genomes to assess the total number of distinct orthogroups with a significantly associated gene. The accumulation curves for 1-19 genomes were produced using the “specaccum” function in the R package *vegan* (Oksanen et al. 2011). We fitted an Arrhenius nonlinear model to the gene accumulation curve to visualize the distribution using the “random” and “fitspecaccum” commands. UpSetR package (Lex et al. 2014) was used to visualize the number of significantly associated genes identified by the multiple reference-based GWAS and K-mer GWAS. All other figures were generated using the R packages *qqman* (Turner, 2014) and *ggplot2* v.3.1.0 (Wickham, 2016).

## Data availability

All genome sequences are available from the NCBI Sequence Read Archive (BioProject accessions PRJNA327615, PRJNA596434, and PRJNA178194).

## Author contributions

AD and DC conceived the research. AD conducted experiments, performed data analyses, and wrote the manuscript with DC. BAM provided funding. All co-authors edited the manuscript.

## Acknowledgments

Emile Gluck-Thaler provided helpful comments on a previous version of the manuscript. This work was supported by the Swiss Federal Office for Agriculture (BLW) in the framework of the NAP-PGREL Project Nr. 627000640.

