## Supplementary Figure for "Combined reference-free and multi-reference approaches uncover cryptic variation underlying rapid adaptation in microbial pathogens"

### Supplementary information

#### Supplementary Figures

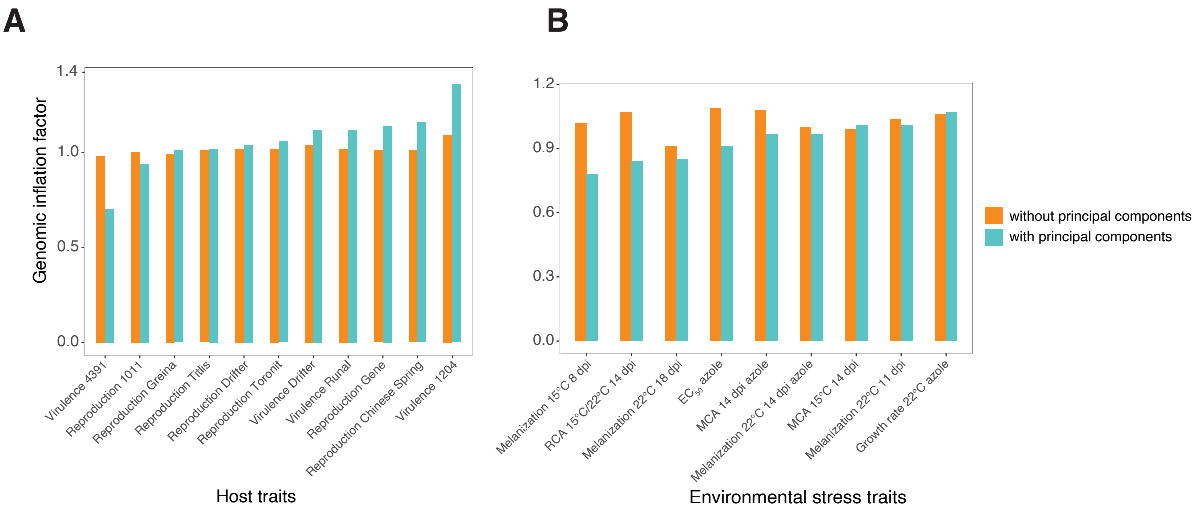


**Supplementary Figure S1.** Genomic inflation factor estimated from genome wide association mapping using principal components as covariates and without principal components in **(A)** host-related traits *i.e.* pathogen virulence (percentage of the leaf surface covered by necrotic lesions) and reproduction (pycnidia density within lesions) and **(B)** environmental stress related traits. Pathogen virulence and reproduction were measured on 12 genetically diverse wheat lines.

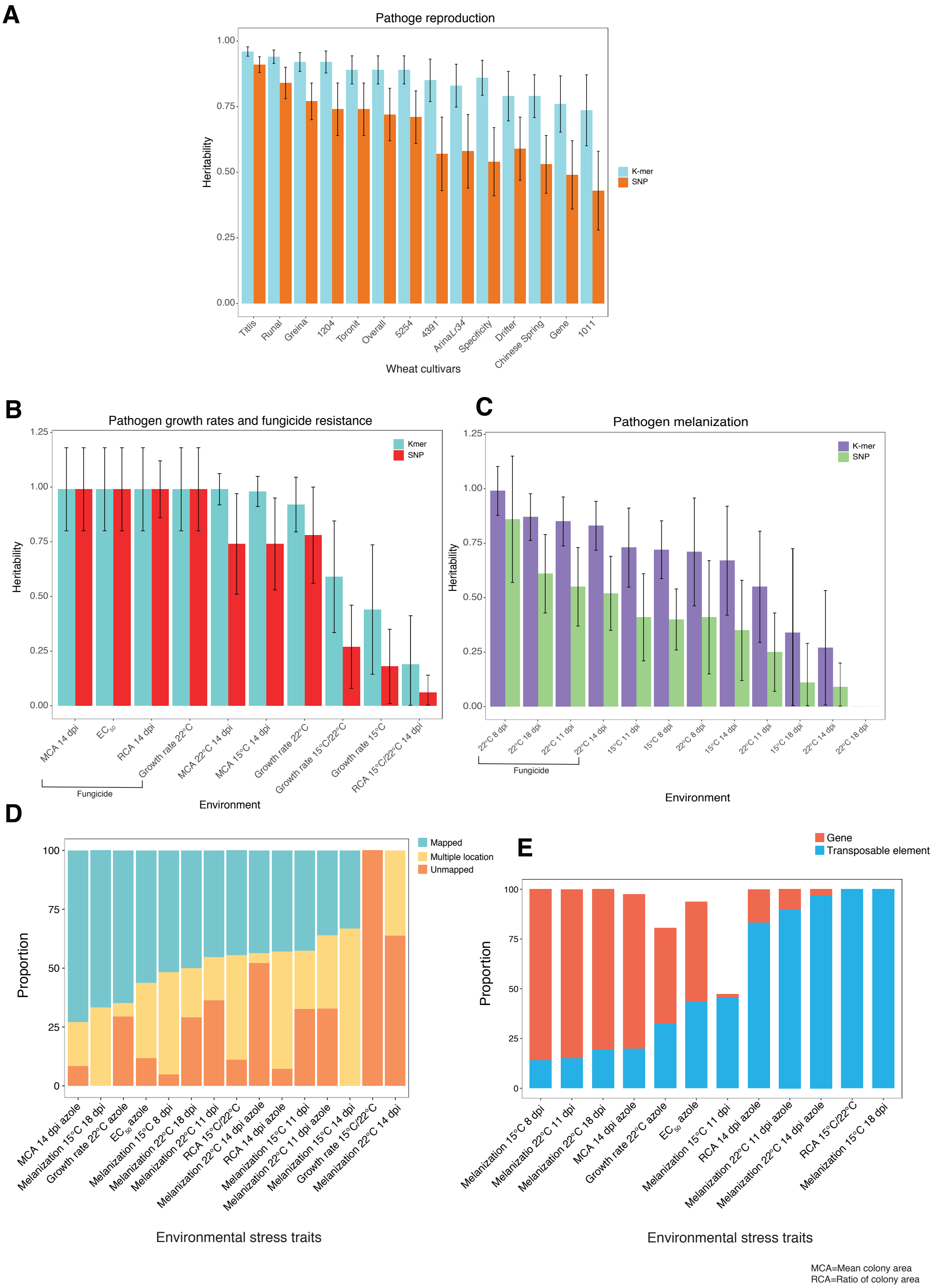


**Supplementary Figure S2.** Comparison of heritability estimates based on SNPs (for the reference genome IPO323) and K-mers in **(A)** pathogen reproduction (pycnidia density within lesions), **(B)** pathogen growth rate and fungicide resistance, **(C)** pathogen melanization. Pathogen reproduction were measured on 12 genetically diverse wheat lines. Overall reproduction represent the average value of reproduction measured on 12 genetically diverse wheat lines. Reproduction specificity was estimated based on the adjusted coefficient of variation of mean reproduction across 12 genetically diverse wheat lines. Higher specificity suggests affinity to certain hosts for maximizing reproductive fitness. Both SNP-based and K-mer-based heritability were estimated by following a genome-based restricted maximum likelihood (GREML) approach. Standard errors are indicated by error bars. **(D)** Alignment of significantly associated K-mers against the reference genome (IPO323) show the proportion of K-mers having a unique mapping position, multiple locations, or no unambiguous mapping position in environmental stress-related traits. **(E)** Proportion of significant K-mers with a unique mapping position in the reference genome either tagging a gene or a transposable element in environmental stress-related traits.

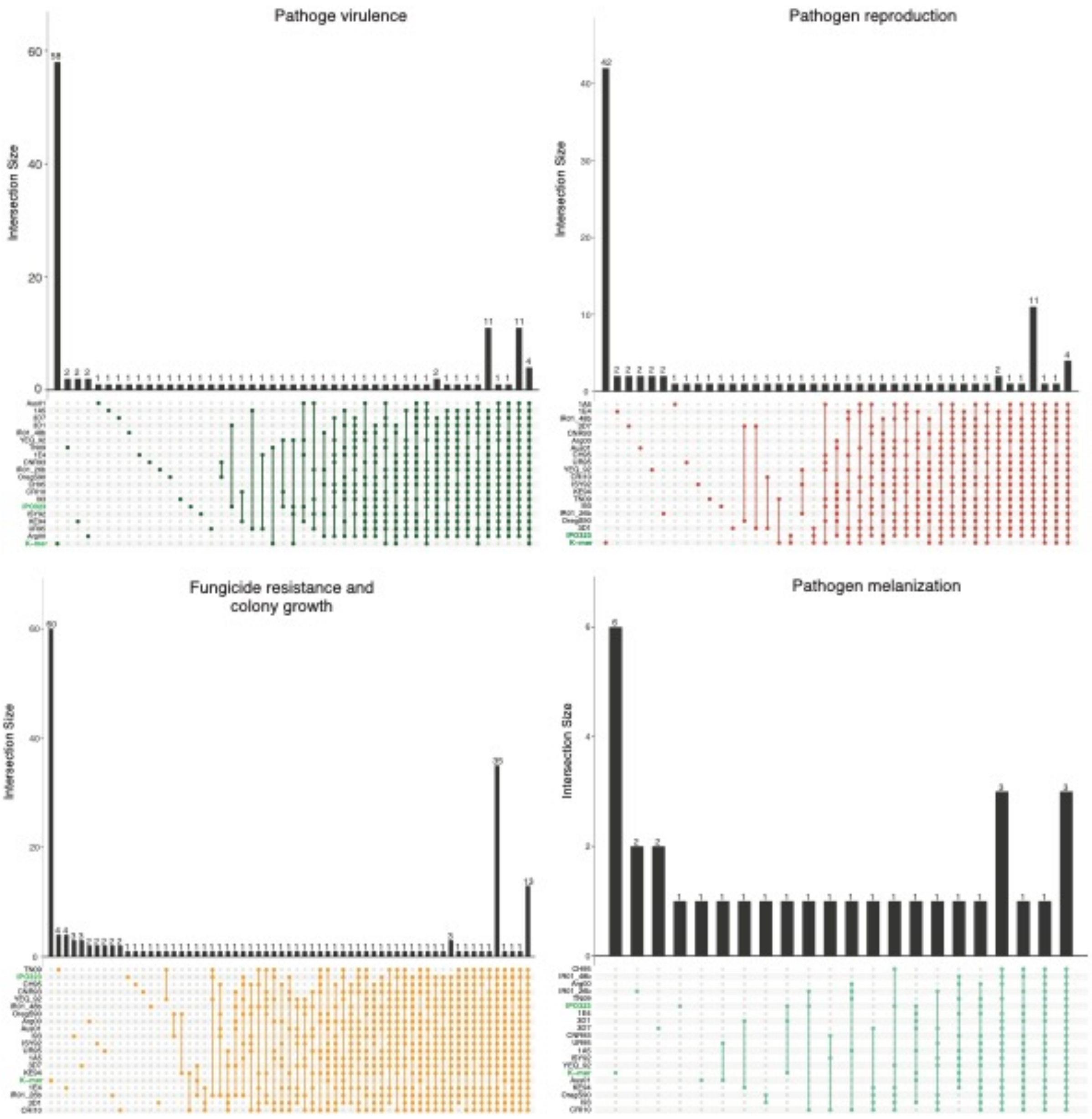


**Supplementary Figure S3.** Upset plots showing the number of genes identified in pathogen virulence (percentage of the leaf surface covered by necrotic lesions), reproduction (pycnidia density within lesions), fungicide resistance and pathogen melanization that are unique or shared among 19 reference genomes in *Zymoseptoria tritici* and the K-mer approach. The intersection size is the number of genes and the black dots on the matrix represent whether the genes are shared or unique to different reference genomes and the K-mer approach. For example, the first vertical bar in each graph shows the number of genes that are uniquely identified by the K-mer GWAS, while the last vertical bar demonstrates the number of genes that are commonly identified by all the reference-based and K-mer GWAS. Pathogen virulence and reproduction were measured on 12 genetically diverse wheat lines.

#### Supplementary Tables

(see separate Excel file)

**Supplementary Table S1.** Raw phenotypic data for virulence (percentage of the leaf surface covered by necrotic lesions) and reproduction (pycnidia density within lesions) on 12 genetically diverse wheat lines from 145 *Zymoseptoria tritici* isolates.

**Supplementary Table S2.** Raw phenotypic data for mean colony area per plate and mean grey value per plate measured in different temperatures and in presence/absence of fungicide from 130 *Zymoseptoria tritici* isolates. "NA" indicates that no data were obtained due to no colony growth or contamination.

**Supplementary Table S3.** Description of 145 *Zymoseptoria tritici* isolates with their corresponding sampling location, year and NCBI SRR Run ID for the whole genome sequence data used in this study.

**Supplementary Table S4.** Number of single nucleotide polymorphisms (SNPs) called on 19 reference genomes at 5% minor allele frequency and 80% genotyping rate for 145 *Zymoseptoria tritici* isolates.

**Supplementary Table S5.** Number of significant SNP associations above the 5% Bonferroni significance threshold for 20 traits comprising pathogen virulence, reproduction and environmental stress mapped in 19 reference genome SNP datasets.

**Supplementary Table S6.** Summary statistics of genome-wide SNPs passing the Bonferroni significance threshold of 5% for specific traits identified 19 reference genome SNP datasets. SNPs are ordered according to the smallest *P*-value.

**Supplementary Table S7.** List of genes with their predicted protein functions in close proximity (< 1 kb) to significant SNPs above the Bonferroni significance threshold (alpha = 0.05) across 19 reference genome SNP datasets for different traits of *Zymoseptoria tritici* isolates.

**Supplementary Table S8.** List of genes with their predicted protein functions in close proximity (< 1 kb) to significant K-mers above the permutation-based significance threshold (5%) for different traits of *Zymoseptoria tritici* isolates..
